# Golden Gate Assembly of Aerobic and Anaerobic Microbial Bioreporters

**DOI:** 10.1101/2021.07.23.453609

**Authors:** Aaron J. Hinz, Benjamin Stenzler, Alexandre J. Poulain

**Author notes:** **Correspondence:** Dr. Alexandre Poulain.

## Abstract

Microbial bioreporters provide direct insight into cellular processes by producing a quantifiable signal dictated by reporter gene expression. The core of a bioreporter is a genetic circuit in which a reporter gene (or operon) is fused to promoter and regulatory sequences that govern its expression. In this study, we develop a system for constructing novel *Escherichia coli* bioreporters based on Golden Gate assembly, a synthetic biology approach for the rapid and seamless fusion of DNA fragments. Gene circuits are generated by fusing promoter and reporter sequences encoding yellow fluorescent protein, mCherry, bacterial luciferase, and an anaerobically active flavin-based fluorescent protein. We address a barrier to the implementation of Golden Gate assembly by designing a series of compatible destination vectors that can accommodate the assemblies. We validate the approach by measuring the activity of constitutive bioreporters and mercury and arsenic biosensors in quantitative exposure assays. We also demonstrate anaerobic quantification of mercury and arsenic in biosensors that produce flavin-based fluorescent protein, highlighting the expanding range of redox conditions that can be examined by microbial bioreporters.

**IMPORTANCE:** Microbial bioreporters are versatile genetic tools with wide-ranging applications, particularly in the field of environmental toxicology. For example, biosensors that produce a signal output in the presence of a specific analyte offer less costly alternatives to analytical methods for the detection of environmental toxins such as mercury and arsenic. Biosensors of specific toxins can also be used to test hypotheses regarding mechanisms of uptake, toxicity, and biotransformation. In this study, we develop an assembly platform that uses a synthetic biology technique to streamline construction of novel *Escherichia coli* bioreporters that produce fluorescent or luminescent signals. We validate the approach by synthesizing and testing an array of bioreporters, including arsenic and mercury biosensors, that produce signal outputs in environments ranging from aerobic to highly reduced anaerobic growth conditions.

## INTRODUCTION

Bioreporters are living host organisms that produce a quantifiable signal in response to the prevailing environmental conditions (1). By providing access to the inner workings of living cells, microbial bioreporters have fundamentally contributed to the knowledge of microbial genetics, physiology, and ecology (2–4). In the field of environmental toxicology, bioreporters offer a simple and cost-effective alternative to analytical methods for quantifying toxic chemicals, while conveying crucial biological insights about toxin bioavailability and cellular stress responses (4).

Bioreporters can be categorized based on whether they produce constitutive or inducible signal outputs, also referred to as “light off” or “light on” biosensors. The constitutive “light off” bioreporters emit a continuous signal and are widely used for fluorescent cell labeling and toxicological screening where the signal intensity decreases with the presence of a toxic response (4, 5). In contrast, inducible “light on” bioreporters produce a signal induced by the intracellular presence of a specific analyte or stressor (1, 5). Inducible biosensors are particularly effective at detecting toxic metals and metalloids that bind with high specificity to regulatory proteins (4), and can be far more sensitive, biologically relevant, and practical than other analytical detection methods (6). Because signal production depends on the bioaccessibility (e.g., access to the cell wall) and bioavailability (e.g., transport across the cell wall) of the analyte (2, 7), biosensors allow researchers to investigate genetic and environmental factors that modulate uptake and transformation of toxins in living cells (8, 9). In addition, biosensors enable the rapid high-throughput screening of environmental samples prior to more costly and in-depth analyses, and can provide an affordable and portable alternative for rapid decision making when analytical support is limited (e.g., in developing countries or remote areas) (4, 10).

The functionality of a bioreporter is determined by promoter sequences that activate expression of a reporter gene (or operon) that produces a quantifiable signal (e.g., green fluorescent protein, luciferase, etc.). Inducible bioreporters can additionally encode regulatory factors that control reporter gene expression in response to a substrate (e.g., the mercury-binding MerR regulator) (11). Bioreporter construction thus involves fusing reporter genes to a promoter and regulatory sequences to create synthetic genetic circuits that are maintained in the host organism on replicating plasmids or integrated into the chromosome (2). The process, however, is often laborious and challenging to scale up, generally relying on traditional molecular cloning methods that involve sequential restriction enzyme digestion, DNA cleanup, and ligation reactions. Synthetic biology methods have emerged as promising alternatives for applications such as DNA assembly (BioBricks, Gibson assembly, Golden Gate assembly, USER fusion, yeast recombination cloning), gene synthesis, and genome editing (MAGE, CRISPR) (12–17). These methods have the potential to vastly increase the efficiency and scale of DNA manipulations but can be challenging for researchers to implement. A major barrier to the implementation of synthetic biology procedures is the requirement of compatible destination vectors, which often have a limited scope of application. To take full advantage of novel cloning methods, it is necessary to increase the diversity, functionality, and accessibility of destination vectors (18, 19).

In this study, we use Golden Gate assembly, a synthetic biology method that produces seamless DNA fusions (20, 21), to build genetic circuits for bioreporters. The approach relies on Type IIS restriction endonucleases, which in contrast to traditional endonucleases, cleave DNA outside of enzyme recognition sequences. The absence of recognition sequence “scars” in the ligated DNA allows for seamless fusion of DNA fragments in “one-pot” cloning reactions that can be used to rapidly construct multi-fragment assemblies, while eliminating time-consuming intermediate steps (20). We present here a streamlined bioreporter construction platform based on the interchangeable assembly of promoter and reporter gene inserts with novel Golden Gate destination vectors. We validated the method by constructing constitutive bioreporters and analyte biosensors that express fluorescent proteins, bacterial luciferase, and a flavin-based fluorescent protein that is active in both aerobic and anaerobic growth environments. Using a common *E. coli* host strain, we measured the signal strength of constitutive reporters, the detection dynamics of mercury and arsenic biosensors, and optimized anaerobic detection of bioreporters that produce flavin-based fluorescent protein.

## RESULTS AND DISCUSSION

### Novel Golden Gate destination vectors

We constructed novel destination vectors that accommodate the assembly of promoter-reporter gene fusions in “one-pot” Golden Gate assembly reactions involving the Type IIS restriction enzyme BsaI (Fig. 1). Each vector encodes a DNA assembly site, consisting of a *lacZ* cassette bounded by two unique BsaI restriction sites. Digestion of the destination vector produces 4-base pair overhangs (“sticky ends”) available for ligation with the compatible overhangs of digested insert DNA. Notably, the overhang sequences were defined to specify the position and orientation of the ligated fragments within the final construct.

**Figure 1.**
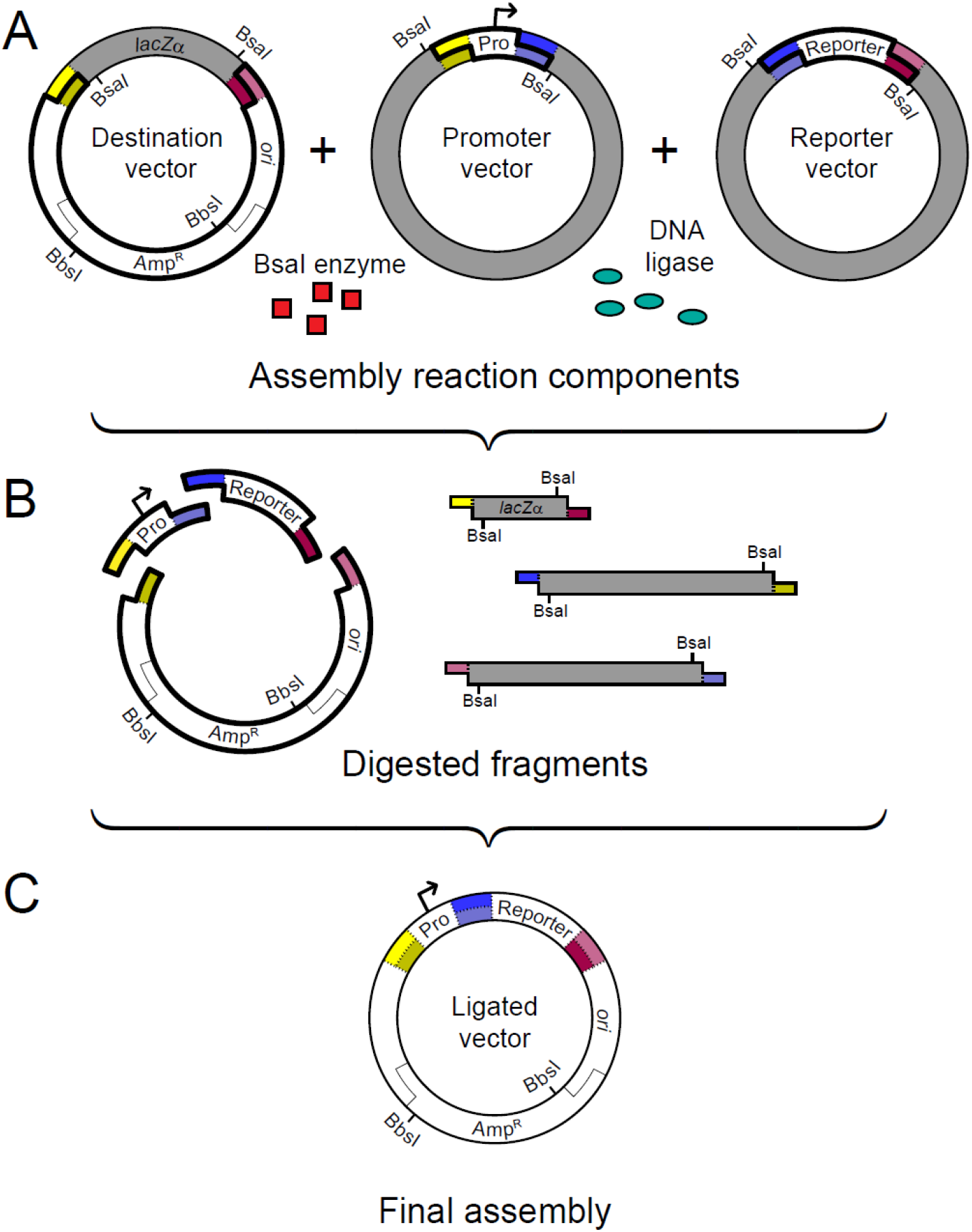
Golden Gate assembly of gene fusions into destination vectors. The diagram illustrates a typical “one pot” assembly reaction that produces a promoter-reporter gene fusion in a Golden Gate destination vector. (A) Reaction components include the destination vector, insert storage vectors, BsaI restriction enzyme, and DNA ligase. The positions of recognition sequences (labelled “BsaI”) and cleavage sites (colored boxes) are positioned such that recognition sequences are absent from fragments intended for the final assembly (outlined in bold). Thus, the enzyme cleaves away from *lacZ*α sequences in the destination vector, and towards the inserts in storage vectors. (B) The sequences of each four-base pair overhang are complementary to those of their intended ligation partner. BsaI recognition sequences are confined to digested fragments excluded from the final construct (gray fill). (C) The assembled vector is resistant to further digestion and lacks *lacZ*α sequences, permitting blue-white screening of transformants. Further modifications to the vector backbone (e.g., substitution of the ampicillin resistance (Amp^R^) cassette) can be performed at the indicated BbsI restriction sites.

Individual destination vectors vary in functions affecting host range, mobilization, plasmid curing, and chromosomal integration (Table 1 and Fig. S1). Basic replicative vectors encode a high-copy number *E. coli* origin of replication (pMB1) (22), an ampicillin resistance cassette, and transcriptional terminators positioned to insulate transcriptional activity originating from insert sequences. Conjugative vectors contain an *oriT* sequence for mobilization into hosts (23, 24). Counter-selectable vectors carry the *sacB* gene for plasmid curing on sucrose media. Chromosomal modification vectors contain elements for site-specific transposition and two-step chromosomal allelic replacement (25–27).

**Table 1.**
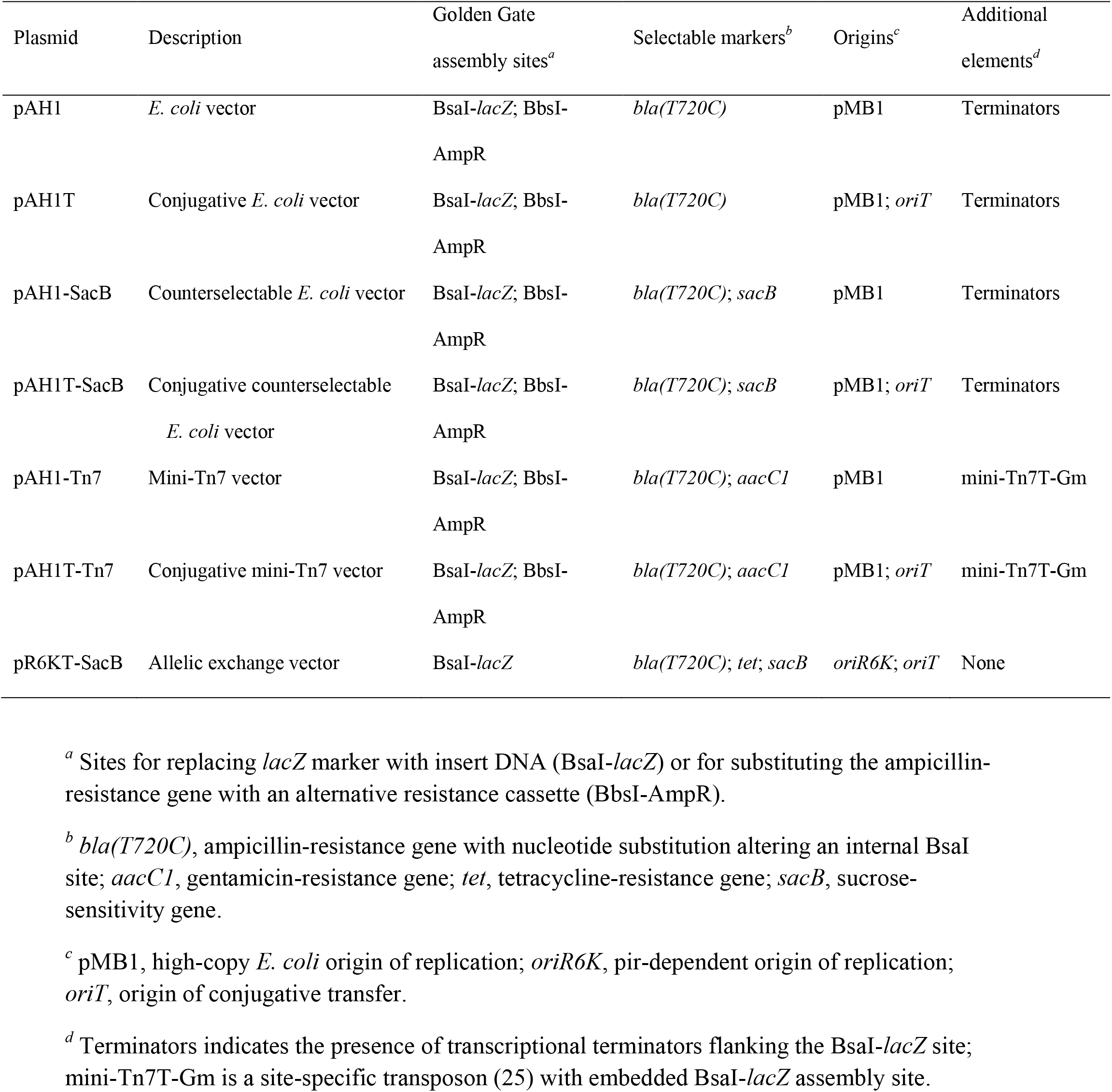
Properties of Golden Gate destination vectors.

A secondary Golden Gate assembly site was included in destination vectors to enable replacement of the ampicillin resistance gene with alternative selectable markers. The cloning site features restriction sites for the Type IIS enzyme BbsI and operates independently of the primary BsaI assembly site. Marker exchange at the BbsI site allows vector backbones to be modified to extend their application to hosts with intrinsic resistance to ampicillin. Furthermore, Golden Gate assembly of multiple DNA inserts at this site can increase versatility by enabling the fusion of selectable marker genes to plasmid elements such as alternative replication origins.

### Combinatorial assembly of bioreporters from promoter and reporter gene inserts

The Golden Gate approach is well-suited for bioreporter construction since it enables the seamless fusion of promoter and reporter inserts. Our design strategy required the modification of insert sequences for directed assembly at the BsaI cloning site of destination vectors. The promoter inserts include three constitutive promoters of varying strength and three inducible promoters responsive to arabinose, mercury, or arsenic (Fig. 2A). The reporter inserts encode yellow fluorescent protein (YFP), mCherry, a flavin-based fluorescent protein (PpFbFP), and bacterial luciferase. We also modified four antibiotic resistance gene inserts for selectable marker replacement at the secondary BbsI cloning site of destination vectors.

**Figure 2.**
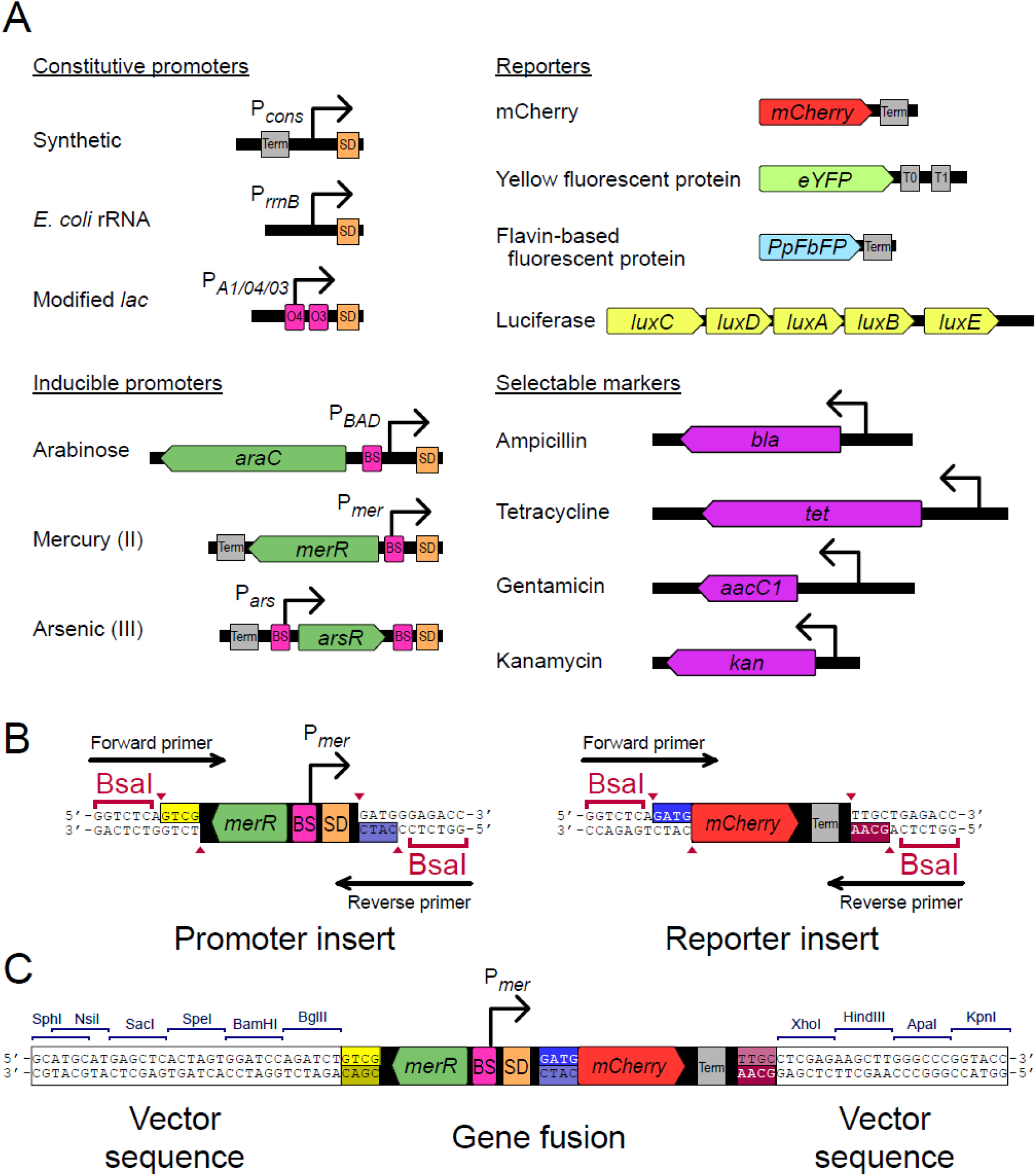
Interchangeable promoter and reporter inserts for bioreporter constructions. (A) Constitutive promoters include a medium-strength synthetic promoter (P*_cons_*) derived from a combinatorial sequence library (47, 48), an *E. coli* ribosomal RNA operon promoter (P*_rrnB_*), and a strong modified T7 promoter with O4 and O3 *lac* operator sites (P*_A1/04/03_*) (28). Inducible promoters responsive to arabinose, mercury, and arsenic are linked to genes encoding their respective analyte-binding transcriptional regulators (8, 9, 49). The locations of regulator binding sites (BS), transcriptional terminators (Term), and Shine-Dalgarno ribosomal binding sites (SD) are indicated. Reporter inserts encode oxygen-dependent (YFP and mCherry) and flavin-dependent (PpFbFP) fluorescent proteins and the bacterial luciferase operon (32, 50, 51). Selectable marker inserts include antibiotic resistance cassettes that can be cloned into the BbsI cloning site of destination vectors. (B) Promoter and reporter inserts were amplified from template sources using primers encoding 5’ sequence extensions encoding the BsaI recognition sequence (5’-GGTCTCN-3’) followed by the unique four-base pair cleavage site sequence (yellow, blue, and purple boxes). The “sticky end” sequences at the promoter-reporter fusion junction include the start codon (ATG) of the reporter gene. (C) Following assembly in Golden Gate destination vectors, gene fusions can be subcloned into alternative vectors using standard restriction sites.

The key modification to insert sequences was the precise addition of BsaI recognition sites and overhang sequences (Fig. 2B). These sequences were routinely added to insert sequences during PCR-amplification via 5’ sequence extensions to the primers. If necessary, existing Type IIS restriction sites within the insert sequences were altered by site-directed mutagenesis (see Methods). Inserts can be included in assembly reactions either as linear PCR products or from storage vectors, as depicted in Fig. 1. During the assembly reaction, defined overhang sequences on the destination vector ligate to complementary overhangs on the promoter and reporter inserts (Fig. 2C). Ligation between the promoter and reporter inserts occurs at a position overlapping the start codon of the reporter gene, with the Shine-Dalgarno (SD) sequences for ribosomal binding located in the promoter insert. The consistent use of four-base pair BsaI overhang sites permits the combinatorial assembly of gene fusions between any pair of promoter and reporter inserts, and novel promoters or reporters can be incorporated by simply adding appropriate BsaI restriction site sequences during insert amplification.

### Signal production of constitutive fluorescent and luminescent bioreporters

The continuous signal emitted by constitutive bioreporters can be used to label cells for microscopy or flow cytometry applications or serve as a proxy of cellular health for general toxicity bioreporters. The specific application of a bioreporter depends on type and level of signal production, which can be impacted by the choice of promoter, reporter, and base vector. We compared the signal outputs of constitutive bioreporters expressing fluorescent or luminescent reporter genes paired with each of three constitutive promoters (P*_rrnB_*, P*_cons_*, and P*_A1/04/03_*). Signal output was quantified in cell suspension assays and is presented as induction rates for fluorescent reporters (Fig. 3A-C), which were stable over the first four hours of the assay, or peak luminescence for luciferase reporters (Fig. 3D), which exhibited non-linear induction (Fig. S3). Although differences in scaling prevent direct comparisons of values measured for different reporter species, the overall patterns illustrate how the type of destination vector, promoter, and reporter species impacts signal production. Among the fluorescent bioreporters, the P*_A1/04/03_* promoter generally produced higher induction rates than the P*_cons_* and P*_rrnB_* promoters (Fig. 3A-C), consistent with the strong activity recognized for this promoter (28). This promoter ranking was observed for mCherry fusions constructed in multiple Golden Gate destination vectors (pAH1 and pAH1T; see Fig. S1) and two established high-copy number vectors (pUC19 and pUCP19) (22, 29), although the choice of destination vector affected the magnitude of signal production (Fig. 3A). Among the flavin-based fluorescent protein (PpFbFP) bioreporters, only the construct with the strong P*_A1/04/03_* promoter produced a higher signal than the “empty vector” control, suggesting that the threshold of PpFbFP detection requires greater gene expression relative to the other fluorescent reporters. Interestingly, the P*_A1/04/03_*-luciferase bioreporter was genetically unstable, likely because of the energy burden imposed by the chemiluminescence reaction, while the lower strength promoters produced a robust signal (Fig. 3D). Overall, these results indicate that the constitutive fluorescent reporters produce reliable and linear signal induction rates, and that the luciferase reporters exhibit higher sensitivity, but with non-linear induction rates.

**Figure 3.**
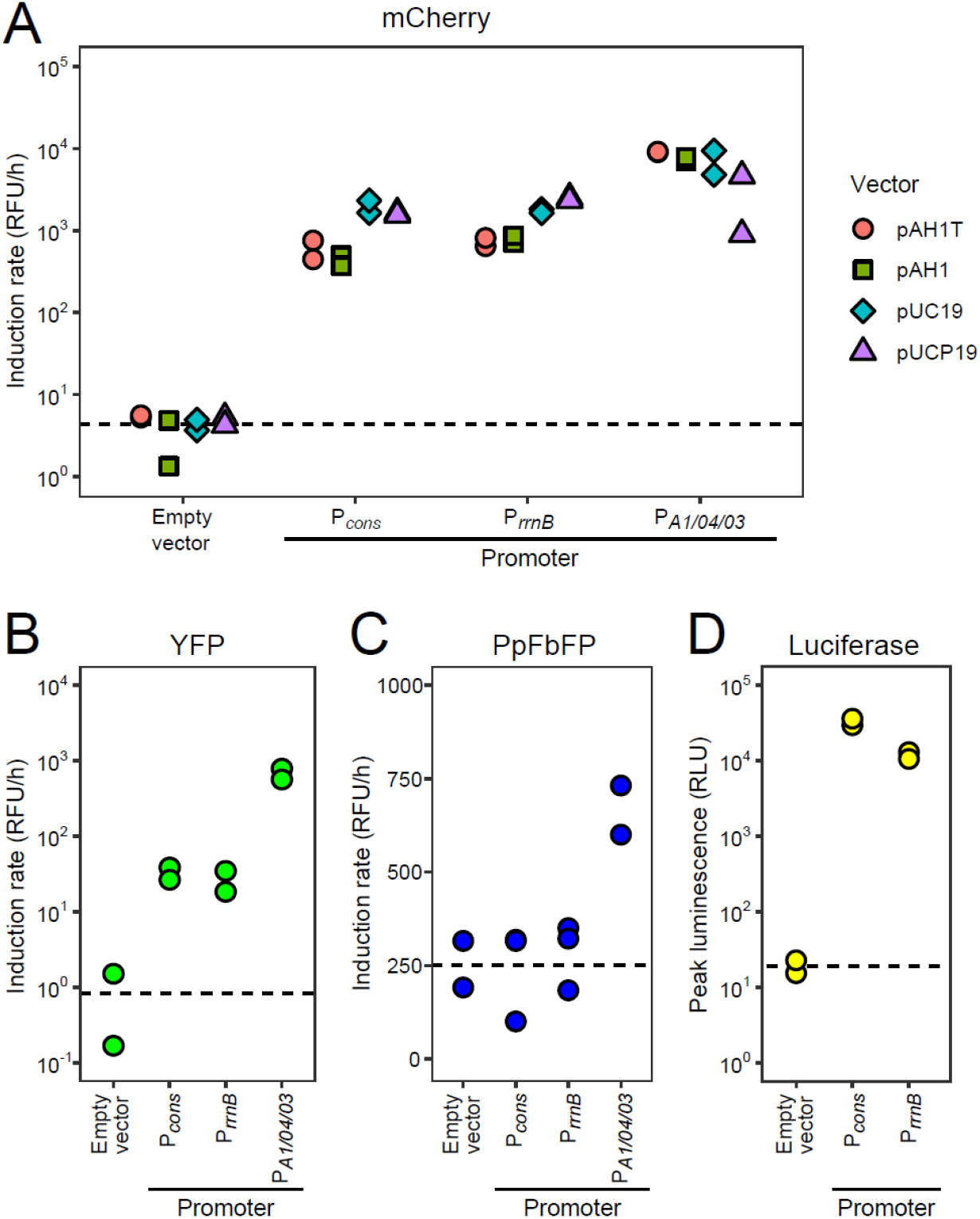
Signal output of constitutive bioreporters expressing mCherry, YFP, PpFbFP, and luciferase. Signal output was quantified in aerobic cell suspension assays of *E. coli* NEB5-α harbouring constitutive (A) mCherry, (B) YFP, (C) PpFbFP, or (D) bacterial luciferase gene fusions. mCherry fusions were assembled in two Golden Gate destination vectors (pAH1T and pAH1) and subcloned into two standard high-copy vectors (pUC19 and pUCP19). The remaining bioreporters were constructed with the pAH1T vector. Signal output is presented as induction rates (relative fluorescence units (RFU) per hour) or as peak luminescence (relative luminescence units (RLU)). The mean values of three technical triplicates are plotted for at least two independently prepared suspension assays for each bioreporter. Dashed lines indicate the mean background signal for “empty vector” cultures. The relative units should only be compared within each reporter type, since scaling is specific to the filter set combination used for signal detection. The mCherry induction rates were transformed by adding a constant of 4 RFU per hour to allow log-scale plotting of negative values obtained for two “empty vector” controls.

### Inducible biosensors for quantification of bioavailable mercury and arsenic

We next evaluated biosensors that produce a reporter signal in response to the binding of intracellular mercury (Hg) or arsenic (As) species to cognate transcriptional repressors. We constructed four Hg-biosensors containing mCherry, YFP, luciferase, and PpFbFP reporters, and quantified their signal responses following exposure to a gradient of Hg(II) concentrations (Fig. 4). Each mercury biosensor produced a dose-dependent response; however, the sensitivity and dynamics of the response depended on the reporter gene. The mCherry and YFP biosensors exhibited linear dose responses between the limit of detection and the maximum concentration tested (1 nM to 20 nM (Fig. 4A and 4B)). In contrast, the luciferase and PpFbFP biosensors were more sensitive near the lower detection limit and exhibited signal plateauing at the higher tested concentrations (Fig. 4C and 4D). The luciferase reporter produced greater variation between biological replicates, however.

**Figure 4.**
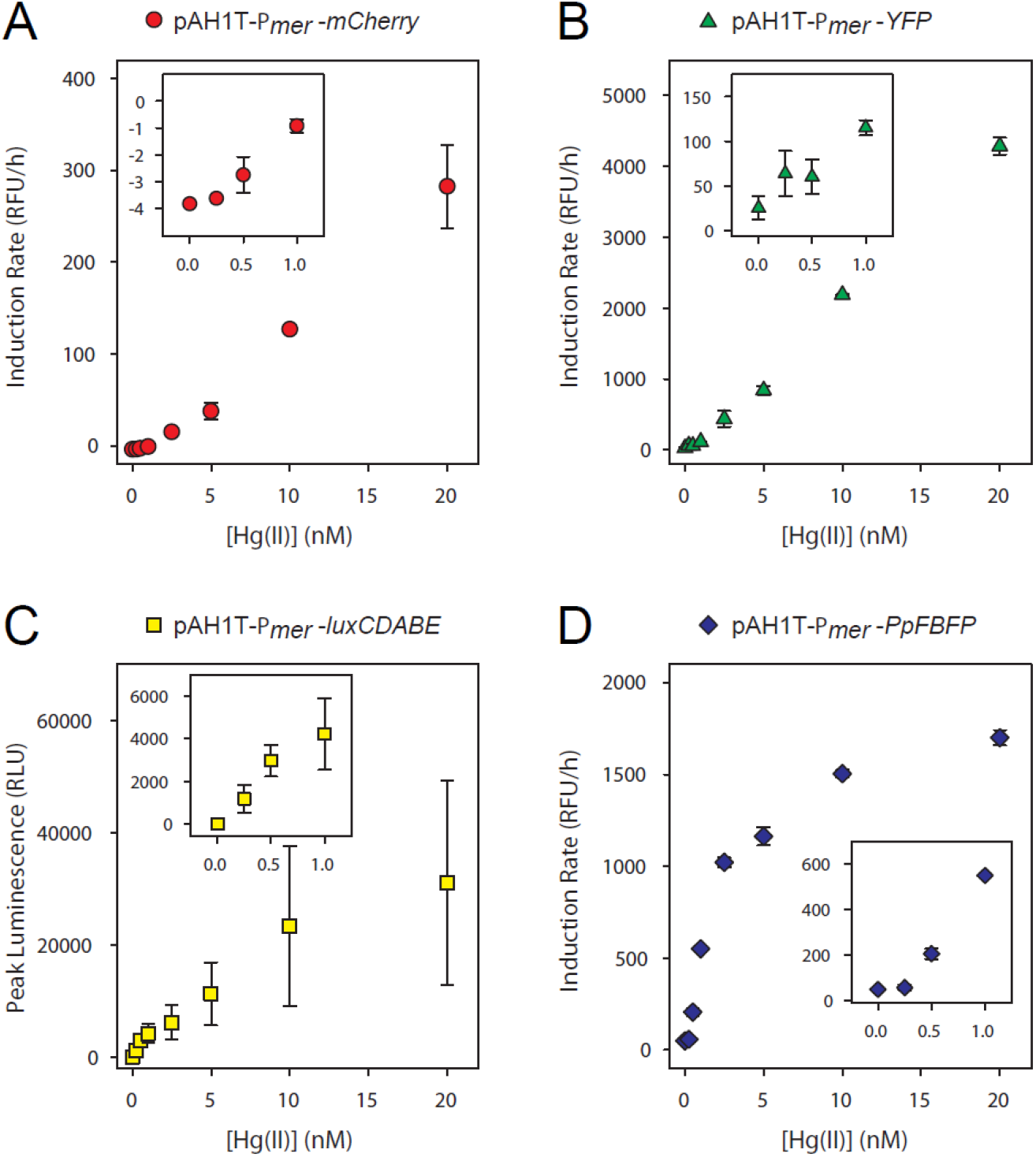
Mercury-inducible biosensors. Mercury-inducible biosensors were exposed to HgSO_4_ concentrations ranging from 0.25 nM to 20 nM. The signal induction rate at each Hg(II) concentration is presented for (A) mCherry, (B) YFP, (C) luciferase, and (D) PpFbFP reporters. The mean induction rate (or peak luminescence) and standard deviation for two biological replicates (assayed in triplicate) is presented for each reporter. Insets were rescaled to display the responses at the lowest tested mercury concentrations.

We constructed a similar set of arsenic biosensors and exposed them to As(V), which is reduced to As(III) under the conditions of the exposure assay (9). The mCherry, YFP, and luciferase reporters exhibited broadly similar response curves between 1 nM and 250 nM with some differences in the sensitivity, level of plateauing, and variation between replicates (Fig. 5A-C). Surprisingly, the PpFbFP arsenic biosensor had a much higher limit of detection (250 nM) than the other reporters, resulting in a major shift in its quantitative range (Fig. 5D). This outcome contrasts with that of the mercury-PpFbFP biosensor, which exhibited a detection limit similar to the alternative mercury biosensors (Fig. 4D). Taken together, these results validate the combinatorial approach to building biosensors from regulatory and reporter insert fragments. Each combination yielded a biosensor producing a distinct dose-response curve, with the choice of reporter gene affecting the sensitivity and the dynamic range of the response.

**Figure 5.**
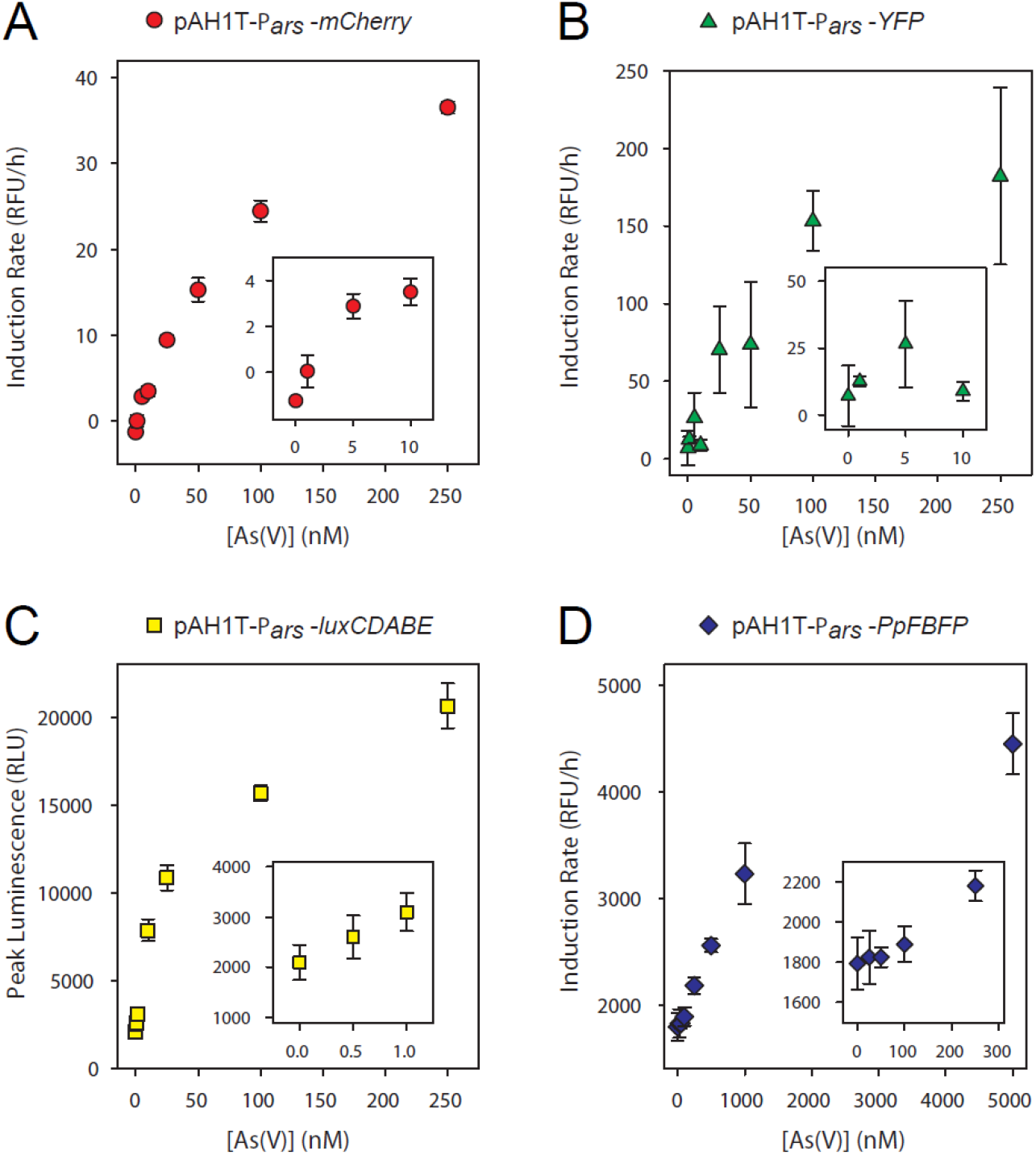
Arsenic-inducible biosensors. Arsenic-inducible biosensors were exposed to As(V) concentrations ranging from 0.5 nM to 5000 nM. The signal induction rate at each As(V) concentration is presented for (A) mCherry, (B) YFP, (C) luciferase, and (D) PpFbFP reporters. The mean induction rate (or peak luminescence) and standard deviation for two biological replicates (assayed in triplicate) is presented for each reporter. Insets were rescaled to display the responses at the lowest tested arsenic concentrations.

### Anaerobic detection of flavin-based fluorescent protein bioreporters

Signal production by light-emitting bioreporters usually requires oxygen for either chromophore maturation or as a reactant in the luciferase reaction (30, 31). Flavin-based fluorescent proteins (FbFPs), on the other hand, produce an oxygen-independent fluorescent signal that is triggered by binding between the fluorescent protein and flavin mononucleotide molecules (30, 32). Therefore, FbFPs can be used to investigate microbial processes under anaerobic conditions in which fluorescent or luminescent bioreporters are typically inactive (8). Signal production by *E. coli* FbFP bioreporters has been demonstrated in cultures undergoing anaerobic respiration with nitrate supplied as terminal electron acceptor (30, 8, 33) but detection in alternative anaerobic growth conditions is less well studied. We tested whether the signal was detectable in cultures undergoing anaerobic respiration with fumarate, a highly reduced terminal electron acceptor. In general, preconditioning the *E. coli* cells through several growth cycles was necessary for reliable anaerobic growth and signal production (see Methods). We found that bioreporters expressing PpFbFP from the strong P*_A1/04/03_* promoter produced a fluorescent signal during both nitrate and fumarate respiration (Fig. S4). Interestingly, the nature of the carbon source was critical for signal detection during fumarate respiration. Cultures supplied with glucose, glycerol, and pyruvate produced a signal, while those supplied with acetate, ethanol, or lactate did not (Fig. S4B), suggesting that both the nature of the carbon source and its redox state are important to consider when designing biosensor assays.

The successful detection of PpFbFP signal from cells undergoing fumarate respiration expands the range of redox conditions for fluorescent bioreporters into highly reduced anoxic conditions. We next tested whether the PpFbFP biosensors were capable of mercury or arsenic quantification during fumarate respiration, an energy metabolism employed by microorganisms that carry out environmental mercury methylation (34–36). We found that the anaerobic dose-response to mercury and arsenic exhibited comparable detection dynamics as the aerobic response (Fig. 6). Overall, these results show that *E. coli* flavin-based biosensors are capable of quantifying analytes under anaerobic growth conditions beyond nitrate respiration, offering opportunities to investigate metal bioavailability and transformation under ecologically relevant culture conditions.

**Figure 6.**
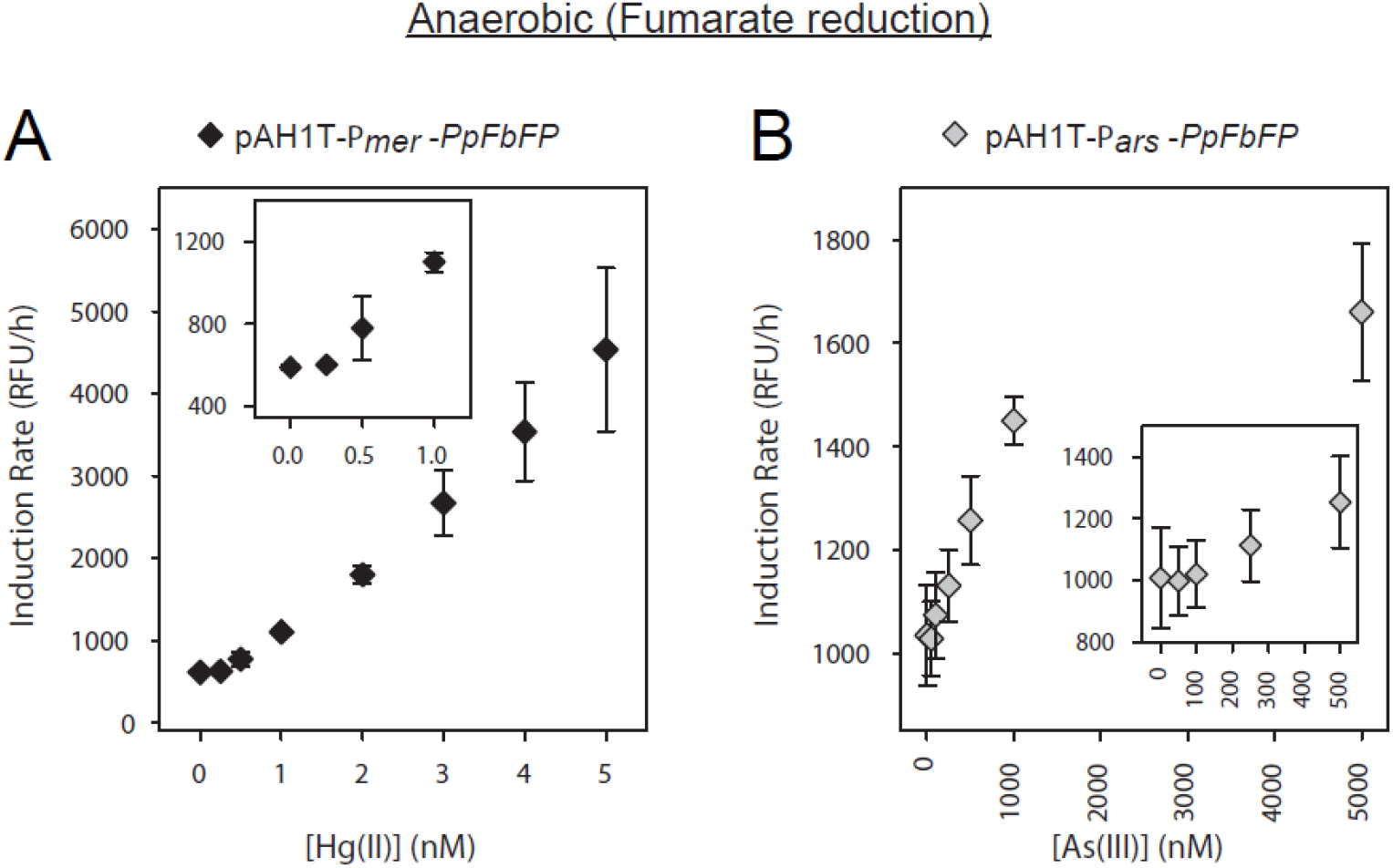
Anaerobic quantification of mercury and arsenic during fumarate respiration. PpFbFP biosensors were exposed to (A) mercury or (B) arsenic in anaerobic conditions with fumarate supplied as terminal electron acceptor. The mean induction rate and standard deviation for two biological replicates (assayed in triplicate) is presented for each reporter as relative fluorescent units (RFU) per hour. Insets were rescaled to display the responses at the lowest analyte concentrations.

## CONCLUSIONS

The aim of this work was to develop a system that applies Golden Gate assembly to molecular cloning applications with a specific focus on bioreporter construction. The advantages of the Golden Gate method are considerable (20, 21). Ligation of multiple DNA segments in a single assembly reaction streamlines cloning procedures and enables the rapid combinatorial assembly of simple or complex gene circuits. Novel insert sequences can be routinely incorporated by simply encoding sequence extensions in PCR primers (see Fig. 3B and Methods), or by custom gene synthesis. We validated our approach by quantifying the outputs of constitutive and inducible bioreporters that produce fluorescent or luminescent signals, including anaerobic FbFP bioreporters that are active during both nitrate and fumarate respiration.

Although the bioreporters we evaluated include only those involving *E. coli* replicative plasmids, we present additional vectors that provide a wide scope of functionalities for bioreporter design and chromosomal modification (Table 1 and Fig. S1). Vectors elements supporting conjugative transfer (*oriT*) (23) and counterselection (*sacB*) (37) can be exploited in mobilizable and curable *E. coli* vectors (38). For microbial species that do not support pMB1-based plasmid replication, the vectors can be utilized for chromosomal allelic replacement (*sacB* vectors) (26, 39) or site-specific transposition (mini-Tn7 vectors) (25, 40, 27). In particular, the pR6KT-SacB allelic replacement vector, which has a restrictive R6K origin (41), has been routinely used to generate chromosomal insertions, deletions, or single nucleotide polymorphisms in *E. coli* (unpublished work) and multiple *Pseudomonas* species (27, 42–44).

Finally, modification of the vector backbone sequences can further extend their host ranges. Alternative replicons can be inserted on restriction sites located in the vector backbones to construct shuttle vectors capable of replication in more diverse hosts. In addition, a second Golden Gate assembly site was engineered specifically to facilitate the substitution of alternative selectable marker sequences. This site can accommodate multi-fragment assemblies and can be exploited to customize destination vectors to specific microbes. For example, host-specific promoters can be fused to selectable marker genes and alternative replicon sequences in assembly reactions. Overall, these properties built into the destination vector design provide a flexible framework for increasing the scope of microbial bioreporter and molecular genetic applications.

## MATERIALS AND METHODS

### Media and molecular biology reagents

*E. coli* cultures were routinely grown at 37 °C in lysogeny broth (LB): 10 g bacto-tryptone, 5 g yeast extract, and 10 g NaCl per liter (45). LB media was supplemented when appropriate with 14 g/l agar (Oxoid), 100-200 μg/ml ampicillin, 30 μg/ml kanamycin, 10 μg/ml gentamicin, 10 μg/ml tetracycline, or 40 μg/ml X-Gal (5-Bromo-4-chloro-3-indolyl β-D-galactopyranoside). Antibiotics and X-Gal were purchased from Sigma-Aldrich.

For the bioreporter suspension assays, cells were grown in LB and Growth Minimal Media (GMM) and exposed in exposure medium. GMM was prepared as described (8). Exposure medium consisted of 40 mM MOPS free acid, 7 mM Sodium Beta-Glycerophosphate, 1 mM (NH_4_)_2_SO_4_, 10 mM Glucose, 5 mM NaOH, and 10-15 mM KOH (to adjust the pH to 7) in Milli-Q water. In anaerobic assays, the exposure media contained 5 mM NaNO_3_ (Nitrate reduction) or 10 mM Sodium Fumarate (Fumarate reduction).

Fumarate-GMM plates were made by autoclaving 191 ml of Milli-Q water with 3.75 g agar in a sealable borosilicate bottle. The media was bubbled with N_2_ while cooling to roughly 60 °C, and the following reagents were quickly added: 43.5 ml of 5X M9 salts, 250 µl of Trace 1 and 2 (as described) (8), 250 µl of 10 mM EDTA, 125 µl of 2 M MgSO_4_, 10 ml of 1 M Sodium Fumarate, 15 µl of 0.225 M FeSO_4_ in 0.1 M H_2_SO_4_, 0.5 ml of 500 mM Thiamine HCl, 1.5 mL of 75 mM Leucine/Isoleucine/Valine solution, 5 ml of 1 M Glucose, 250 µl of 100 mg/ml Ampicillin. Once reagents were added, the bottle was sealed very tightly and cycled into the anaerobic chamber. Plates were poured anaerobically and used the day they were prepared.

Custom oligonucleotides were purchased from Invitrogen (Thermo Fisher Scientific). Polymerase chain reactions (PCR) were prepared with Phusion High-Fidelity DNA Polymerase (Thermo Fisher Scientific). PCR products were purified with the Wizard SV Gel and PCR Cleanup System (Promega). Plasmid DNA was isolated from bacterial cultures with the QIAprep Spin Miniprep Kit (Qiagen). Restriction endonucleases and T4 DNA ligase were purchased from New England Biolabs. High-fidelity versions of BsaI and BbsI enzymes were used in Golden Gate assembly reactions.

### Golden Gate assembly and molecular cloning methods

Golden Gate assembly reactions were prepared as previously described (20, 21). Approximately equimolar amounts (∼40 fmol) of destination vector and insert storage vectors were mixed with 1 μl of 10X T4 ligase buffer, 0.5 μl (200 units) of T4 ligase, and 0.5 μl (10 units) of Type IIS enzyme (e.g., BsaI) in a total volume of 10 μl. Reactions were incubated for 2 h at 37 °C, 5 min at 50 °C, and 5 min at 80 °C. Chemically competent *E. coli* DH5α λpir were transformed with 2 μl of the assembly reactions by the Inoue method (45), and the cells were incubated for 2 h at 37 °C to allow expression of resistance genes. Cells were plated on LB agar supplemented with 100 µg/ml ampicillin and X-Gal. Inserts were validated by colony PCR of white colonies or Sanger sequencing (Genome Quebec, McGill University). Confirmed plasmids were isolated and transformed into chemically competent *E. coli* NEB5-alpha cells (New England BioLabs) for quantitative exposure assays.

Traditional restriction enzyme cloning was used in the construction of destination and storage vectors. For these reactions, vector and insert DNA were digested separately for 2 h at 37 °C, purified by spin column, and ligated for 16 h at 16 °C before transformation into *E. coli*. TOPO cloning of purified Phusion PCR products into the pCRBluntII-TOPO vector (Thermo Fisher Scientific) was performed according to the manufacturer’s protocol.

### Design of Golden Gate assembly sites

Two Golden Gate assembly sites were designed for ligation of reporter gene fusions (BsaI-*lacZ* site) and selectable marker cassettes (BbsI-AmpR site) to destination vectors. The BsaI-*lacZ* site was derived by PCR amplification of *lacZ*α sequences from pUC19 (22) with primers introducing the restriction site sequences (F2-pUC19-BsaI and R2-pUC19-BsaI; Table S2). The PCR amplicon was cloned into the BglII and SpeI sites of allelic exchange vector pAH385 (a derivative of pAH79) (42, 46) to form pR6KT-SacB (Table S1). In subsequent destination vector designs, the BsaI-*lacZ* site was situated within the mini-Tn7 polylinker (25) (Fig. S2), to provide restriction site options for subcloning inserts into alternative vectors. The BsaI-*lacZ* site was transferred to the mini-Tn7 polylinker by amplifying the BsaI-*lacZ* cloning site from pR6KT-SacB with primers F-MCS and R-MCS (Table S2) and ligating the BbsI-digested product at the BamHI and XhoI sites of the polylinker.

The Golden Gate marker exchange site (BbsI-AmpR) includes a beta-lactamase (*bla*) cassette flanked by inverted BbsI recognition sequences that produces restriction overhangs compatible with selectable marker inserts. The site was created by amplifying the *bla(T720C)* allele (containing a mutation altering an internal BsaI site) with primers F-marker and R-marker (Table S2 and Table S3) and incorporating the amplicon into vector backbones during destination vector assembly.

### Destination vector constructions

Novel destination vectors were constructed by Golden Gate assembly of multiple PCR amplicons encoding vector functions (e.g., replication origin, selectable markers, and transcriptional terminators) into closed ligation loops. Details on the properties of source and constructed vectors, primer sequences, and amplicons used for vector assemblies are found in Tables S1-S3. Intermediate destination vectors containing at minimum the pMB1 replicon and BbsI-AmpR marker exchange site were generated by BsaI-mediated assembly of PCR amplicons listed in Table S3. In a second step, the BsaI-*lacZ* assembly site and surrounding mini-Tn7T polylinker were cloned into the intermediate vectors (see Table S1 for specifics). New functions can be added to or replace existing functions on vector backbones by exploiting standard restriction sites engineered at the fusion junctions between the assembled vector segments. For example, the counterselectable vectors pAH1-SacB and pAH1T-SacB were derived by excising the pMB1 origin from their parent vectors and substituting an amplicon containing pMB1 and *sacB* sequences.

### Promoter, reporter, and selectable marker inserts

Inserts compatible with the Golden Gate destination vectors were created by PCR amplification of target sequences from source vectors (Table S1-S3). Terminal sequence tags of the primer sequences include: 1) six non-specific base pairs at the termini for efficient cleavage near the ends of DNA fragments; 2) an optional standard restriction site for cloning insert segments into long-term storage vectors; 3) a Type IIS recognition sequence (BsaI for bioreporter assembly or BbsI for marker exchange); 4) a defined four-base pair overhang sequence that directs ligation between digested fragments; and 5) an optional standard restriction site for subcloning assembled gene fusions. Consistency in the four-base pair overhang sequences allows for the pairing of any combination of promoter and reporter inserts. Novel inserts can thus be integrated into the assembly platform by using the primer sequences listed in Table S2 as guides. Notably, the fusion junction between promoter and reporter inserts overlaps with the start codon of the reporter gene. Therefore, new promoter inserts must include appropriately distanced ribosome binding sites, and new reporter inserts must be in-frame with the start codon located within the fusion junction.

To avoid interference in assembly reactions, all secondary BsaI (5’-GGTCTC-3’) and BbsI (5’-GAAGAC-3’) recognition sequences in destination vector and insert sequences were altered. We introduced synonymous mutations at secondary sites using a previously described site-directed mutagenesis method that introduces mutation at ligation junctions during Golden Gate assembly of PCR products (27). Derivatives of pUC19 and pAH79 vectors were constructed that carry a synonymous mutation that alters the internal BsaI site of the beta-lactamase gene: *bla(T720C)*. In addition, the following synonymous nucleotide substitutions were introduced in insert gene sequences: *luxC(A1312C)*, *tetA(C654G)*, and *mCherry(G456A)*. All amplicons carrying promoter, reporter, and selectable marker sequences were cloned into one or more of the following vectors for long-term storage: pCR-BluntII-TOPO (Invitrogen), pUIC3, pUC19, pEX18Gm, and pEX18Tc (22, 39, 46).

### Growth for bioreporter suspension assays

*E. coli* strains were grown at 37 °C and selected with 200 μg/ml ampicillin (unless otherwise stated) in preparation for suspension assays, with no antibiotic used in the suspension assay. For aerobic assays, a single colony was selected from a plate and grown in LB broth over day. Cultures were transferred (1% inoculum) from LB to GMM and grown overnight. The next morning, cultures were transferred (10% inoculum) into fresh GMM and grown to an OD_600_ of 0.6. Cultures were resuspended twice in exposure media (10,000 RCF for 90 seconds). The cultures were ready for a 1/20 dilution in exposure media for the bioreporter suspension assay.

Anaerobic suspension assays were carried out under both nitrate and fumarate reducing conditions for bioreporters expressing flavin-based fluorescent protein. All anaerobic manipulations of cell cultures were performed in a Coy Vinyl Anaerobic chamber (98% N_2_, 1.5% H_2_), and anaerobic growth was performed in anaerobic balch tubes. Before growth for the suspension assay, cultures were acclimated to grow in Anaerobic GMM supplemented with 25 mM NaNO_3_ by culturing for 7-10 days with daily 1% transfers from stationary phase, until an OD_600_ > 0.6 was reached. Corresponding cryostocks were made of the acclimated cultures. From cryostock, cultures were plated onto an LB plate with 120 μg/ml ampicillin and grown aerobically. Single colonies were selected and grown aerobically in LB broth overnight. The following morning, the cultures were resuspended in anaerobic GMM (20% inoculum) and grown over the day. The cultures were transferred (1% inoculum) to grow overnight, then transferred (20% inoculum) and grown to an OD_600_ of 0.6. Cultures were resuspended twice in anaerobic exposure media supplemented with 5 mM NaNO_3_ (10,000 RCF for 90 seconds). The cultures were ready for a 1/20 dilution in exposure media for the bioreporter suspension assay.

The FbFP bioreporter suspension assay was optimized for fumarate reduction using a pUC19 construct with the promoter *P_A1/O4/O3_,* which produced the highest anaerobic response on NO_3_^-^ (Fig. S4A). This constitutive bioreporter as well as the analyte-sensing bioreporters were acclimated to grow in anaerobic GMM supplemented with 40 mM Na_2_Fumarate by culturing for 7-10 days with daily 1% transfers from stationary phase, until an OD_600_ > 0.6 was reached. Corresponding cryostocks were made of the acclimated cultures. From cryostock, cultures were plated onto fumarate-GMM plates with 100 μg/ml ampicillin and grown in anaerobic plate jars for several days. A single colony was selected and grown in anaerobic GMM supplemented with 40 mM Na_2_Fumarate. The following day, cultures were transferred (20% inoculum) and grown to an OD_600_ of 0.6. Cultures were resuspended twice (10,000 RCF for 90 seconds) in anaerobic exposure media supplemented with 10 mM Na_2_Fumarate. The cultures were ready for a 1/20 dilution in exposure media for the bioreporter suspension assay.

The bioreporter suspension was optimized for fumarate reduction using a variety of different electron sources that *E. coli* has known metabolism for: acetate, ethanol, glucose, glycerol, lactate and pyruvate (Fig S4B). Glucose was ultimately selected for exposure due to the highest fluorescent induction response. Pyruvate and Glycerol also were able to produce a signal, while acetate, ethanol, lactate, and the no carbon/electron source treatment gave no induction response.

### Bioreporter suspension assays

Suspension assays were performed in exposure medium using 7 ml standard PTFE vials. Hg was prepared as a 7.2 μM HgSO_4_ stock in 0.2 M H_2_SO_4_, the concentration being determined using an MA-3000 Mercury Analyzer. Hg was diluted in exposure medium to a working concentration of 100 nM in PTFE vials before addition to the suspension assay. Arsenic was prepared as a 10 mM Na_2_HAsO_4_ or Na_2_AsO_3_ standards and was diluted in exposure medium in PTFE vials before addition to the suspension assay. Anaerobic exposure medium was supplemented with 5 mM NaNO_3_ or 10 mM Sodium Fumarate. After dilution of the corresponding metal(loid) in the exposure medium, bioreporter cells (see “Growth for bioreporter suspension assays”) were added to 1/20 of the suspension assay volume of 2 ml. The suspension assay was transferred in triplicate from the PTFE vials to a 96-well plate (Black, Clear-Bottom, Nonbinding Surface Microplates) and transferred to a Tecan Infinite F200Pro plate reader (Aerobic) or a Synergy HTX plate reader (Anaerobic).

Readings for fluorescence, luminescence and OD_600_ were taken every five minutes for 20 hours at 37 °C. Fluorescence was read for YFP (Ex: 500 nm, Em: 535nm), mCherry (Ex: 560 nm, Em: 620nm), and PpFbFp (aerobic Ex: 450 nm, Em: 500 nm; anaerobic Ex: 440 nm, Em: 500 nm). No growth was observed in the duration of the exposure assays. Signal induction rates were calculated for inducible and constitutive fluorescent bioreporters for the first 4 hours of exposure or 10 hours for cultures undergoing fumarate respiration. For As inducible luminescence, max initial luminescence peak was quantified within 4-6 hours. For Hg inducible luminescence, a max value was determined within 20 hours.

### Data availability

Sequences of vectors and inserts generated for this study will be deposited to GenBank upon acceptance.

## ACKNOWLEDGEMENTS

We thank Thien-Fah Mah and Rees Kassen for kindly providing genetic resources and members of the Poulain lab, especially Martin Pothier, for helpful insights and discussions. This work was funded by the Natural Sciences and Engineering Research Council of Canada (NSERC).

